# Detection of Prenatal Alcohol Exposure Using Machine Learning Classification of Resting-State Functional Network Connectivity Data

**DOI:** 10.1101/2020.08.14.231357

**Authors:** Carlos I. Rodriguez, Victor Vergara, Suzy Davies, Vince Calhoun, Daniel D. Savage, Derek A. Hamilton

**Affiliations:** The Mind Research Network. 1101 Yale Blvd. NE, Albuquerque, NM, 87106, USA; Tri-Institutional Center for Translational Research in Neuroimaging and Data Science (TReNDS), Georgia State University, Georgia Institute of Technology, and Emory University. 55 Park Pl NE, Atlanta, GA 30303, USA; Dept. of Neurosciences, University of New Mexico School of Medicine. 1 University of New Mexico, Albuquerque, NM, 87131, USA; Department of Psychology, University of New Mexico. 1 University of New Mexico, Albuquerque, NM 87131, USA

**Keywords:** Prenatal Alcohol Exposure, Fetal Alcohol Spectrum Disorder, Machine Learning, Functional Network Connectivity

## Abstract

**Introduction:** Previous work utilizing resting state fMRI to measure functional network connectivity in rodents with moderate prenatal alcohol exposure (PAE) revealed several sex- and region-dependent alterations in FNC implicating FNC as potential biomarker for PAE. Given that FNC is sensitive to neurological and psychiatric conditions in humans, here, we explore the use of previously acquired FNC data and machine learning methods to detect PAE among a sample of rodents exposed to moderate PAE and controls exposed to a saccharin solution.

**Materials & Methods:** We utilized previously acquired fMRI data from 48 adult rats 24 PAE (12 male 12 female) and 24 saccharin exposed (SAC) controls (12 male and 12 female) for classification. The entire data sample was utilized to perform binary classification (SAC or PAE) of FNC data with multiple support vector machine (SVM) kernels and out-of-sample cross-validation to evaluate classification performance.

**Results:** Results revealed accuracy rates of 62.5% for all samples, 58.3% for males, and 79.2% for females using a quadratic SVM kernel to classify moderate PAE from FNC data. In addition, brain networks localized to hippocampal and cortical regions contributed strongly to QSVM classifications.

**Conclusion:** Our results suggest overall modest classification performance of a QSVM to detect moderate PAE from FNC data gathered from adult rats, yet good performance among females. Further developments and refinement of the technique hold promise for the detection of PAE in earlier developmental time periods that potentially offer additional tools for the non-invasive detection of PAE from FNC data.

**IMPACT STATEMENT:** The diagnosis of fetal alcohol spectrum disorders (FASDs) can be challenging in individuals who lack the hallmark facial dysmorphologies associated with heavy prenatal alcohol exposure (PAE). The absence of a diagnosis prevents individuals with PAE from receiving the treatment and services that improves quality of life outcomes. This research explores the use of preclinical functional network connectivity data and machine learning techniques as a novel and non-invasive means of detecting PAE. Our aim is to contribute basic science towards improving diagnostic strategies that can lead to securing timely and appropriate support for individuals with FASD and their caregivers.

## INTRODUCTION

Fetal alcohol spectrum disorder (FASD) is a term that is utilized to encompass a wide range of morphological and neuro-behavioral phenotypes caused by exposure to alcohol during prenatal development (Loock, Conry, Cook, Chudley, & Rosales, 2005; Williams, Smith, & Committee On Substance, 2015). The most severe end of the spectrum is known as Fetal Alcohol Syndrome (FAS) and is linked to heavy prenatal alcohol exposure (PAE) (Lemoine, Harousseau, Borteyru, & Menuet, 1968; Manning & Eugene Hoyme, 2007). Children with FAS exhibit facial dysmorphologies, growth deficits, and numerous impairments in cognitive and behavioral functions related to attention, learning, memory, and motor coordination among others (Connor, Sampson, Bookstein, Barr, & Streissguth, 2000; Jones & Smith, 1973, 1975; Streissguth et al., 1986). Although the most severe, FAS is the least common with an estimated prevalence rate of ~0.1% in the U.S. (May & Gossage, 2001). However, when considering the entire spectrum, estimated prevalence rates of FASD (including FAS) fall between 1.1% and 5.0% of U.S. children, many of which will not display readily identifiable facial dysmorphologies, but may nonetheless exhibit cognitive and behavioral deficits (May et al., 2014; May et al., 2018). Unfortunately, children who do not display the facial dysmorphologies characteristic of FAS, due to lack of early diagnosis, may not receive timely treatment or services, negatively impacting life outcomes related to academic success (Mattson & Riley, 1998), difficulty finding and maintaining meaningful employment, and staying out of trouble with the law (Popova, Stade, Bekmuradov, Lange, & Rehm, 2011).

From the early clinical descriptions of FAS (Jones & Smith, 1973, 1975), research with human participants has been critical for understanding the social, physical, and neuro-behavioral sequelae of PAE (Connor et al., 2000; Streissguth et al., 2004). However, variables such as dose (e.g., high, moderate, low), timing (e.g., 1^st^, 2^nd^ trimester), and pattern of alcohol exposure (e.g., daily vs binge), can be difficult to account for and, for ethical reasons, are impossible to experimentally manipulate in human subjects research (Patten, Fontaine, & Christie, 2014). To overcome these challenges, animal models of FASD have been important for illuminating the underlying neurobiological consequences associated with developmental alcohol exposure.

Given that children are more likely to be exposed to moderate, rather than heavy, levels of prenatal alcohol exposure (May et al., 2018; May & Gossage, 2001), animal research aimed at studying the effects of moderate PAE is extremely valuable because it more closely mimics the pattern of alcohol exposure observed in the human population. Within animal models of PAE, considerable work has been undertaken with the aim of investigating discrete brain areas such as the hippocampus (Gil-Mohapel, Boehme, Kainer, & Christie, 2010; Savage, Becher, de la Torre, & Sutherland, 2002) and cerebellum (Servais et al., 2007). However, higher level cognitive and behavioral functions, including those associated with FASD, involve sophisticated and highly coordinated activity across multiple, rather than single, brain regions (Green et al., 2009). Functional network connectivity (FNC) methods (i.e. functional connectivity between coherent brain networks) offer an important lens that can be leveraged to understand the temporal statistical dependencies (e.g. correlations) of multiple and distant brain networks (Arbabshirani & Calhoun, 2011) following PAE. Functional magnetic resonance imaging (fMRI), a neuroimaging modality employed to non-invasively measure blood-oxygenation-level dependent (BOLD) signals that reflect patterns of neuronal activity (Logothetis, Pauls, Augath, Trinath, & Oeltermann, 2001; Raichle & Mintun, 2006), has been widely utilized to derive measures of FNC (Allen et al., 2011). Group level fMRI data gathered at rest, an experimental condition that lacks externally presented stimuli or behavioral responses (Snyder & Raichle, 2012), can be examined by group independent component analysis (GICA). As a blind source separation algorithm, GICA is a data driven technique that extracts the temporal activation patterns (time courses) of resting state networks (RSNs) where each network may consist of multiple brain regions (Allen et al., 2011; Arbabshirani & Calhoun, 2011; Buckner, Krienen, & Yeo, 2013). FNC assessment consists of correlations between the time-courses of brain networks. Brain dysfunction can then be identified by abnormal correlations (e.g. too high or too low) when comparing FNC across control and experimental treatment conditions.

Previous work from our group applied GICA to resting state fMRI data acquired from adult rodents exposed to moderate levels of PAE that revealed several sex and regionally dependent alterations in functional network connectivity (Rodriguez, Davies, Calhoun, Savage, & Hamilton, 2016) and point to FNC is a potential biomarker for the identification of PAE. In human-subjects research, measures of FNC from fMRI data have also been successfully utilized to detect patients with schizophrenia and mild traumatic brain injury (mTBI) using machine learning algorithms (Rashid et al., 2016; Vergara, Mayer, Damaraju, Kiehl, & Calhoun, 2017). In this study, we explored the use of multiple binary-classification machine learning algorithms and leave-one-out-cross validation (LOOCV) to detect PAE among a mixed sample of FNC data from alcohol- and saccharin-exposed (SAC; control) rodents. Functional neuroimaging data were obtained from our previously published report that characterized the effects of moderate PAE on FNC by utilizing GICA of resting-state fMRI data (Rodriguez, Davies, et al., 2016). The primary goal of this current investigation was to explore the utility of machine learning algorithms a novel and non-invasive means to detect aberrant patterns of FNC in a rodent model of FASD.

## METHODS

### Subjects

Subjects, materials, and procedures were previously reported in separate studies approved by the Institutional Animal Care and Use Committee of the University of New Mexico main campus and Health Sciences Center (Rodriguez, Davies, et al., 2016; Rodriguez, Magcalas, et al., 2016). Briefly, 48 Long-Evans rats (24 SAC and 24 PAE) were generated in a single breeding round designed to prenatally expose rats to either a 5% ethanol (v/v) or 0.066% saccharin solution (Hamilton et al., 2014) for the duration of the entire 21-day gestational period. Following weaning, animals were housed with an age- and weight-matched cagemate from the same prenatal treatment, but different litter, in standard plastic cages with water and food available ad libitum.

At 3-4 months of age, all animals underwent a series of structural- and blood oxygenation level dependent (BOLD) fMRI-scan sequences under isoflurane anesthesia for ~45 min in a 4.7T Bruker Biospin (Billerica, MA) MRI scanner. Functional MRI data were acquired with a 10-minute echo planar imaging acquisition at a temporal resolution (TR) of 2 sec (FOV = 3.84 cm × 3.84 cm, matrix = 64 × 64, TE = 21.3 ms, flip angle = 90°, 27 slices, and slice thickness = 1 mm).

### Image Preprocessing, Group Independent Component Analysis (GICA), and Functional Network Connectivity

Preprocessing, GICA, and FNC methods are described in (Rodriguez, Davies, et al., 2016). To summarize, fMRI data preprocessing included realignment, spatial normalization to the Paxinos & Watson atlas (Schweinhardt, Fransson, Olson, Spenger, & Andersson, 2003), and smoothing with a 0.5 mm full-width-half-maximum (FWHM) Gaussian kernel in Statistical Parametric Mapping 8 (SPM8) (Wellcome Department of Cognitive Neurology, London, UK) running in MATLAB (Mathworks, Inc., Natick, MA) version R2012b. After preprocessing, 40 group-level independent components were extracted utilizing the Infomax algorithm (Bell & Sejnowski, 1995) in the Group ICA of fMRI Toolbox (GIFT, www.trendscenter.org/software/gift) (Calhoun, Adali, Pearlson, & Pekar, 2001). Of the initial 40 components, 17 components were retained based on the exclusion of components localized to white matter tracts or cerebro-spinal fluid and the presence of artifactual features upon visual inspection.

In this study, component time courses were orthogonalized with respect to the following: (1) linear, quadratic, and cubic trends; (2) the six realignment parameters (translation in the x, y, and z directions and rotations about the x, y, z axes); and the 6 realignment parameter derivatives. Time-courses were lowpass filtered with a cutoff at 0.15Hz. Functional network connectivity (FNC) measures were estimated from pairwise correlations between average individual component time-courses for each rat. A total of 136 unique pairwise correlations were calculated for each animal given by the following: 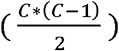 where *C* = 17 (the number of retained components). Thus, the structure of the FNC data utilized for machine learning procedures consisted of 48 correlation matrices. All correlation values were Fisher’s Z transformed for subsequent analyses.

### Machine Learning Procedures

The machine learning methods to classify FNC patterns between PAE and SAC animals was based on work previously described in (Vergara, Mayer, Kiehl, & Calhoun, 2018) and relied on utilizing FNC data from GICA-extracted components. SVM tuning parameters included a least squares solving method, a soft margin parameter of 0.1 and a feature selection threshold of 0.75. Feature selection is implemented by running a two-sample t-test on SAC and PAE groups for each of the 136 FNC values and discarding FNCs with a t-value failing to meet the t = 0.75 threshold. This SVM configuration was used to run five different SVM Kernels: linear, quadratic (QSVM), cubic, radial basis functions (RBF) and multilayer perceptron (MLP) kernels in MATLAB (Mathworks, Inc., Natick, MA) version 2016b to perform binary classification of FNC data at the subject level. Because of the relatively small sample size, leave-one-out cross validation (LOOCV) was chosen to assess classification performance. The LOOCV procedure consisted of isolating one sample for testing and the remainder of the samples for training across multiple iterations as displayed in Figure 1. Statistical significance was assessed by a permutation test approach in which the prenatal condition labels of individual subject’s FNC data were randomized and subsequently subjected to 10,000 replications of QSVM classification and LOOCV with random groups on each replication to establish the null probability distribution of accuracy rates from randomized data (null model). Significance at the p = 0.05 level was estimated from the null distribution. Finally, to address potential differences in sex, the permutation test approach and LOOCV procedures were repeated separately for male and female samples.

**Figure 1.**
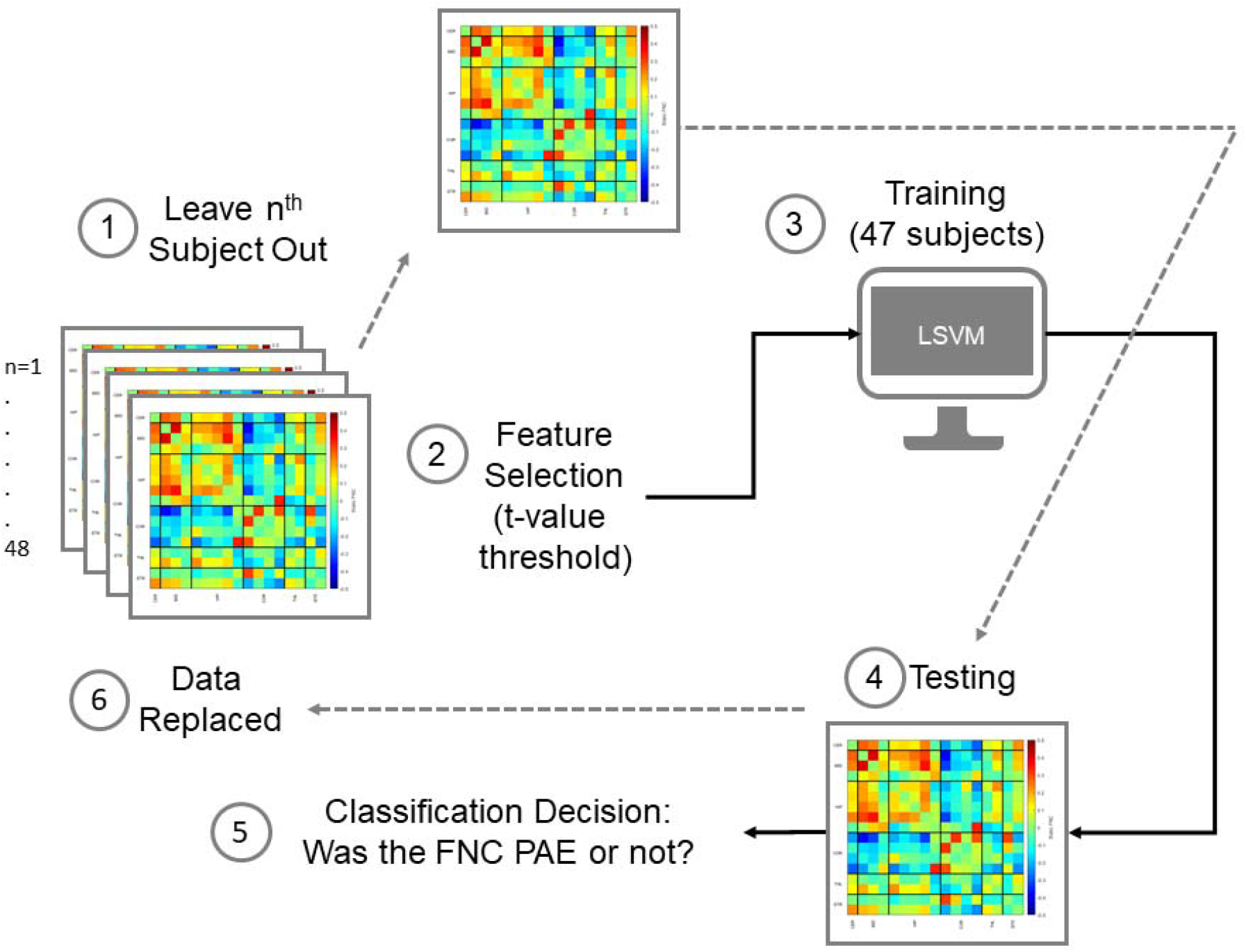
Leave-one-out cross validation schematic illustrates FNC matrices that represent the 136 pairwise correlations of RSNs (blue=negative correlations, red=positive correlations) for each subject (48). For each iteration, the connectivity matrix from the n^th^ subject is left out (1), the remaining 47 matrices underwent feature selection based on a t-value threshold (2), and then utilized for training the SVM (3). The left out, n^th^, subject data is then utilized for measuring (4) performing a classification decision (5). Finally, the nth subject data is replaced (6) and the process reiterates until all 48 classification decisions were gathered. Correct classifications out of 48 comprise SVM accuracy rates.

## RESULTS

Retained components are displayed in Figure 2 and the anatomical location for the peak value of each component, in Paxinos and Watson space, is displayed in Table 1 (Paxinos & Watson, 2004). However, we would like to remind the reader that these components and their location were previously reported in an earlier study and do not represent results from an independent investigation. Components are displayed to aid the reader in localizing the brain regions from which the FNC measures for this study are derived from.

**Figure 2.**
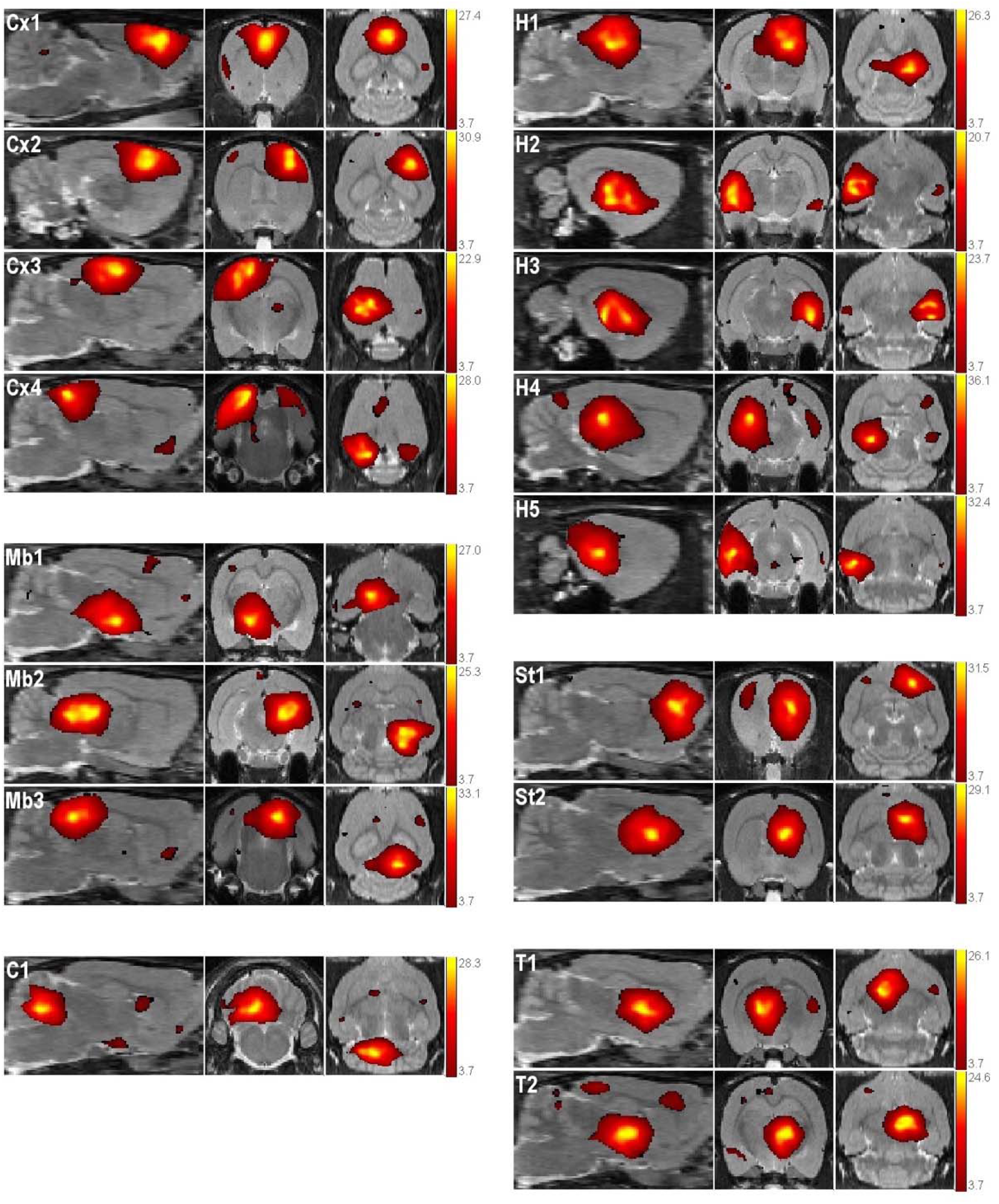
Retained independent components representing RSNs in sagittal, coronal, and axial views. The anatomic location of the peak component t-value determined grouping into cortical (Cx), midbrain (Mb), hippocampal (h), striatal (St), cerebellar (C) and thalamic (T) networks. Reprinted with permission (Rodriguez, Davies, et al., 2016)

**Table 1.**
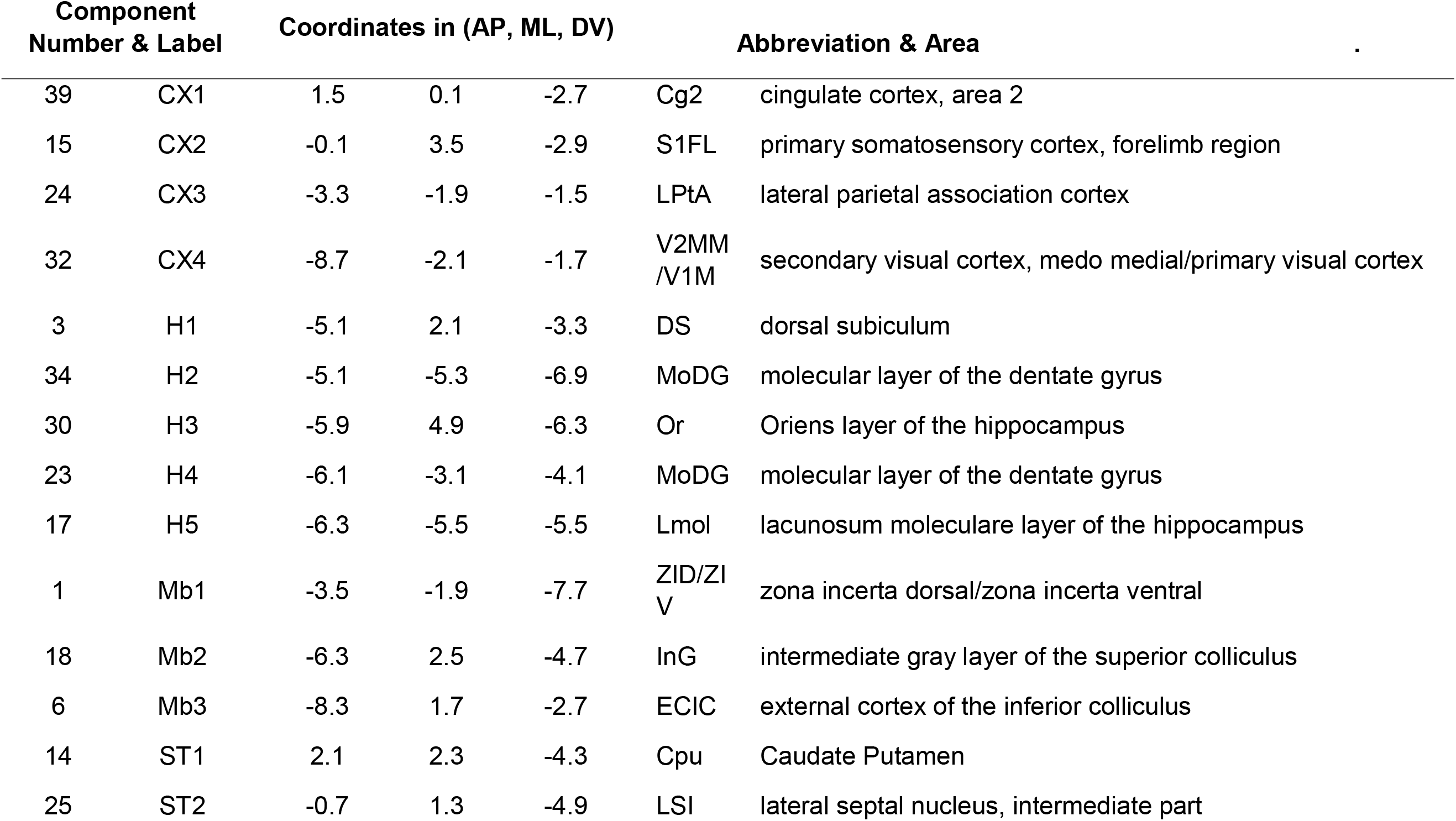
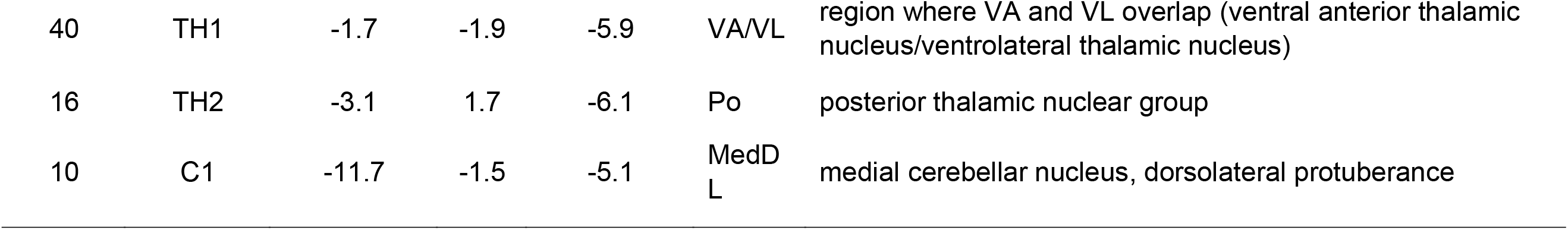
Anatomical locations of extracted components. Components are arranged according to the Paxinos and Watson rat atlas (Paxinos & Watson, 2004) coordinates from anterior to posterior within regional grouping. Cortex (CX), Hippocampus (H), Midbrain (MB), Striatum (ST), and Cerebellum (C). Reprinted with permission (Rodriguez, Davies, et al., 2016).

| Component<br>Number & Label |  | Coordinates in (AP, ML, DV) |  |  | Abbreviation & Area |  | . |
| --- | --- | --- | --- | --- | --- | --- | --- |
| 39 | CX1 | 1.5 | 0.1 | -2.7 | Cg2 | cingulate cortex, area 2 |  |
| 15 | CX2 | -0.1 | 3.5 | -2.9 | S1FL | primary somatosensory cortex, forelimb region |  |
| 24 | CX3 | -3.3 | -1.9 | -1.5 | LPtA | lateral parietal association cortex |  |
| 32 | CX4 | -8.7 | -2.1 | -1.7 | V2MM<br>/V1M | secondary visual cortex, medio medial/primary visual cortex |  |
| 3 | H1 | -5.1 | 2.1 | -3.3 | DS | dorsal subiculum |  |
| 34 | H2 | -5.1 | -5.3 | -6.9 | MoDG | molecular layer of the dentate gyrus |  |
| 30 | H3 | -5.9 | 4.9 | -6.3 | Or | Oriens layer of the hippocampus |  |
| 23 | H4 | -6.1 | -3.1 | -4.1 | MoDG | molecular layer of the dentate gyrus |  |
| 17 | H5 | -6.3 | -5.5 | -5.5 | Lmol | lacunosum moleculare layer of the hippocampus |  |
| 1 | Mb1 | -3.5 | -1.9 | -7.7 | ZID/ZI<br>V | zona incerta dorsal/zona incerta ventral |  |
| 18 | Mb2 | -6.3 | 2.5 | -4.7 | InG | intermediate gray layer of the superior colliculus |  |
| 6 | Mb3 | -8.3 | 1.7 | -2.7 | ECIC | external cortex of the inferior colliculus |  |
| 14 | ST1 | 2.1 | 2.3 | -4.3 | Cpu | Caudate Putamen |  |
| 25 | ST2 | -0.7 | 1.3 | -4.9 | LSI | lateral septal nucleus, intermediate part |  |

|  |  |  |  |  |  |  |
| --- | --- | --- | --- | --- | --- | --- |
| 40 | TH1 | -1.7 | -1.9 | -5.9 | VA/VL | region where VA and VL overlap (ventral anterior thalamic nucleus/ventrolateral thalamic nucleus) |
| 16 | TH2 | -3.1 | 1.7 | -6.1 | Po | posterior thalamic nuclear group |
| 10 | C1 | -11.7 | -1.5 | -5.1 | MedD<br>L | medial cerebellar nucleus, dorsolateral protuberance |

Table 2 displays the accuracy rates of multiple kernels used in SVM binary classification. The quadratic kernel demonstrated the highest classification rates when classifying all (both male and female samples; 62.5%) and female samples only (79.2%). The quadratic and RBF kernels demonstrated the highest accuracy rates for male samples (58.3 %). The lowest accuracy rates were observed for all samples using the RBF kernel (50%), males using the linear kernel (50%) and for females a three-way tie (66.7%) among linear, cubic, and MLP kernels. The quadratic SVM (QSVM) kernel displayed the best overall accuracy rates for discriminating between alcohol- and saccharin exposed animals and was therefore chosen for the remaining series of analyses.

**Table 2.** Classification accuracy rates of different SVM kernels. Support vector machine (SVM), radial basis (RBF), multilayer perceptron (MLP).

| SVM Kernel | All Samples % | Males % | Females % |
| --- | --- | --- | --- |
| Linear | 54.0 | 50.0 | 66.7 |
| Quadratic | 62.5 | 58.3 | 79.2 |
| Cubic | 60.4 | 41.7 | 66.7 |
| RBF | 50.0 | 58.3 | 70.8 |
| MLP | 54.2 | 54.2 | 66.7 |

Figure 3 displays accuracy-rate histograms after the prenatal conditions of FNC data were randomized and subjected to 10,000 iterations of QSVM classification and LOOCV to establish null distributions. Each null distribution was used to estimate the p=0.05 level threshold of statistical significance for accuracy rates of QSVM classification in all (A), female (B), and male samples (C). The significance threshold for all samples was 63%. Therefore, the accuracy rate for QSVM classification (62.5%) was near significance. For females, the significance threshold was estimated at 67% thus the QSVM classification rate for females (79.2%) was statistically significant. Finally, the significance threshold for males was also 67%, rendering the QSVM’s performance for classifying males as not statistically significant.

**Figure 3.**
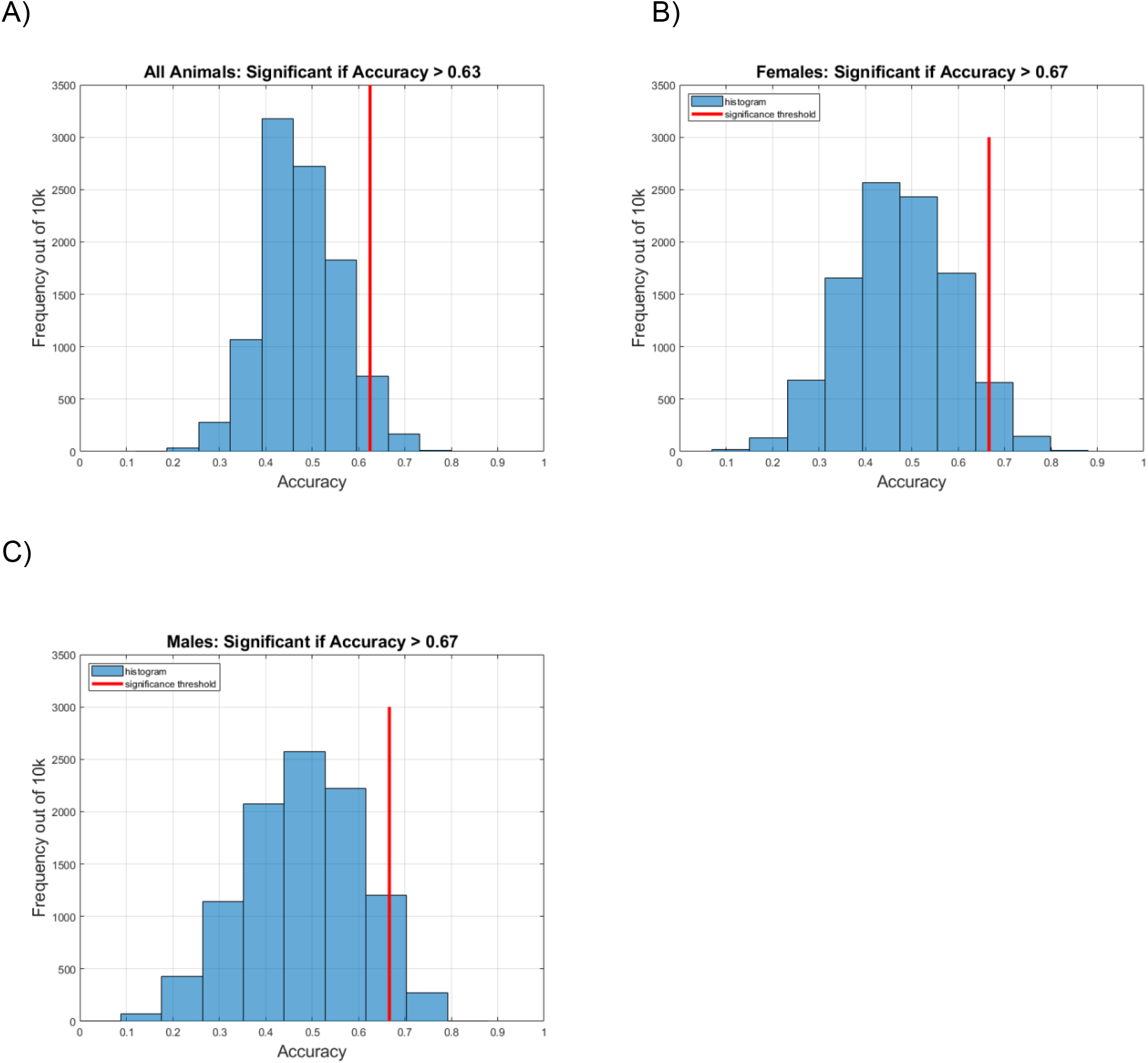
Null-model classification accuracy histograms illustrate the SVM classification accuracies after prenatal condition label randomization and cross validation over 10,000 iterations. The resulting distribution of accuracy rates under the null model provides the basis for calculating a *p*-value for the probability of obtaining an accuracy rate equal to or greater than observed SVM classification accuracy rates for A) All animals, B) Females, and C) Males.

The mean static FNC correlations between all independent components are displayed in Figure 4A. Moderate positive within-network connectivity (triangular regions along the diagonal) is generally observed in hippocampal, midbrain, and thalamic components while negative within network connectivity is exhibited in striatal components. Clear patterns of positive between-network connectivity are observed in midbrain-hippocampal and cerebellar-hippocampal couplings, while negative between-network connectivity is readily observed among cortical-cerebellar, cortical-midbrain, cortical-hippocampal, and cortical-thalamic couplings. Remaining combinations of connectivity display a mix of positive and negative correlations without a readily visually distinguishable pattern. Two-sample t-tests for each pairwise connectivity measure between prenatal conditions (PAE-SAC) are displayed in Figure 4B. Strong differences between conditions can be appreciated in cortical-midbrain, cortical-cortical, cortical - thalamic and cortical-striatal couplings. Separate t-test matrices for females (PAE-SAC) and males (PAE-SAC) are displayed in Figures 4C and 4D respectively. Comparing the female to male matrices reveals many of the differences from the all samples t-tests are driven by female animals. Stronger differences in mean female connectivity can be appreciated in cortical-midbrain, cortical-cortical, and cortical-striatal couplings. Less pronounced differences among male animals are generally observed with reductions in p-value magnitude readily observed in multiple regions with pronounced reductions displayed in cerebellar, cortical, and striatal networks.

**Figure 4.**
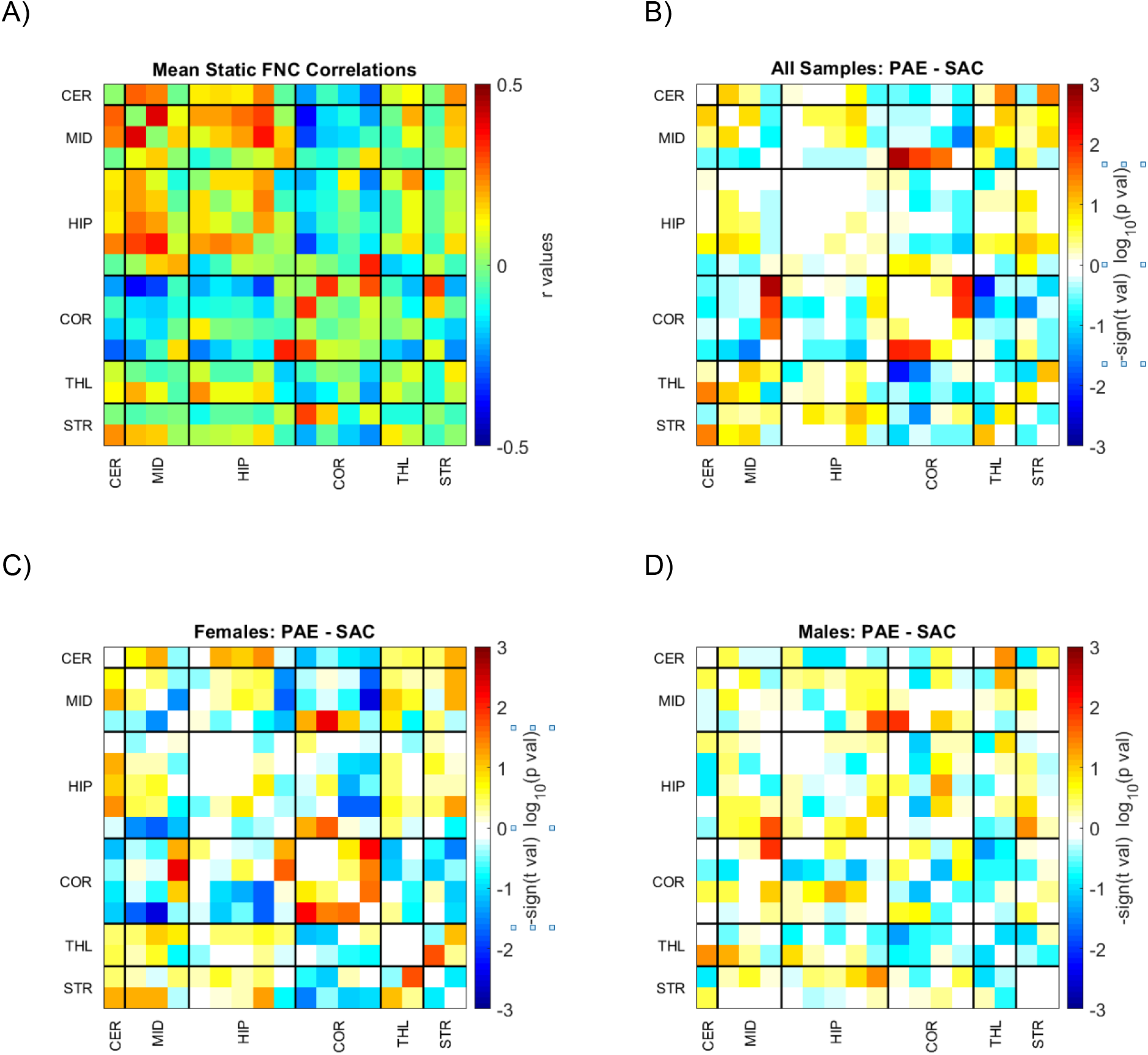
Mean static FNC correlation matrix for all subjects (A). Significance and direction following two-sample t-tests (PAE-SAC) on each pairwise correlation are depicted for all subjects (B), males (C) and females (D) as the -sign(t val)log_10_(p val). Component labels correspond to striatal (St), thalamic (T), cortical (Cx), hippocampal (H), midbrain (Mb), and cerebellar (C) networks.

The QSVM assigns weight to each pairwise correlation used in classification and mean classification weights are displayed in Figure 5 for all samples(A), females (B), and males (C). Weights can be used to explore the contributions of specific component correlations that most strongly impact correct classification decisions.

For all samples, a general pattern of moderately positive weights results from network correlations between cerebellar-hippocampal connectivity. Other moderate positive weights result from couplings in hippocampal-striatal, hippocampal-cortical, and hippocampal-midbrain components, while a strong mean positive weight was found in a hippocampal-thalamic coupling consisting of components with peak activations localized to the ventral-anterior thalamus and the dentate gyrus of the hippocampus. Strong negative weights result from cerebellar-cortical, hippocampal-midbrain, and cortical-striatal couplings.

For males, strong and moderately strong positive weights cluster in cortical-midbrain, cortical-hippocampal, cortical-cortical, cerebellar-hippocampal, and hippocampal-thalamic, midbrain-thalamic, midbrain-striatal, and midbrain-hippocampal connectivity. Strong negative weights are observed between striatal-cortical, cerebellar-cortical, cortical-hippocampal, and midbrain-midbrain, and striatal-thalamic connectivity.

For females, strong and moderately positive weights are observed between cortical-hippocampal, cortical-striatal, striatal-thalamic, cerebellar-hippocampal, and cerebellar-midbrain couplings. Clear patterns of moderately strong negative weights are observed in hippocampal-midbrain, cortical-cortical, cortical-cerebellar, cortical-hippocampal, cortical-cortical, thalamic-hippocampal, striatal-hippocampal, striatal-cortical, and striatal-thalamic couplings.

## DISCUSSION

The motivation for this study was predicated on previous work that demonstrated regional- and sex-dependent differences in FNC patterns following moderate PAE in adult rats. Our goal was to explore the utility of machine learning to perform binary classification from resting state fMRI connectivity data acquired from an animal model of PAE. We found that a quadratic SVM kernel demonstrated the highest classification rates when compared to linear, cubic, RBF, or MLP kernels. QSVM-kernel-based classification resulted in an accuracy rate of 62.5% for all animals, 58.3% for males, and 79.2% for females. To assess statistical significance, we employed a permutation testing approach with 10,000 replications of randomization and LOOCV and found the female classification rate to surpass a *p = 0.05* threshold. Qualitative and quantitative evaluation of FNC data and QSVM weights implicate an overarching theme of several hippocampal and cortical networks in contributing to treatment dependent differences in connectivity and the formation of correct classification decisions by the QSVM.

It is important to note that the blank cells in Figure 5 indicate correlations that did not surpass the feature selection step of the QSVM classification process. When examining these cells, a striking feature of the classification weight data is a considerable reduction in the amount of correlations used in classification of males when compared to females. Thus, a possible reason for the higher classification rate success in female animals may be due to a higher number of correlations that surpassed the t-value threshold that facilitated the classification process. These results also indicate an overall greater degree of differences in FNC across females. Interestingly, in our previous study, we found males displayed more alterations in FNC as a result of moderate PAE (Rodriguez, Davies, et al., 2016). The present work, however, investigated non-linear data features that may be related to the improved classification of PAE in females compared to males. Another key difference for the present report is a more refined time-course pre-processing pipeline which included detrending, regression of realignment parameters, and filtering to account for in-scanner movement and to reduce the potential signal contributions stemming from respiratory processes.

Other resting state fMRI research conducted with rats exposed to prenatal alcohol has also revealed sex-dependent alterations in connectivity. Using a seed-based approach, Tang and colleagues showed baseline sex-dependent differences in functional connectivity among controls and a sex-by-alcohol interaction in cortico-striatal connectivity (Tang et al., 2019). In a subsequent investigation relying on a graph theory approach, the same research group found altered network organization in females, but not males, following PAE (Tang, Xu, Zhu, Gullapalli, & Mooney, 2020). Taken together, these findings point to the existence of sex-related differences in network connectivity in rodents with PAE with underlying mechanisms that are currently poorly understood.

Multiple reports in the literature have shown disrupted resting state functional connectivity following the administration of ketamine, a N-methyl-D-aspartate (NMDA) glutamate receptor antagonist (Grimm et al., 2015; Kraguljac et al., 2017; Motoyama et al., 2019; Mueller et al., 2018) and implicate glutamatergic neurotransmission as a candidate mechanism contributing to PAE-dependent network dysfunction. However, previous work with the same moderate PAE paradigm used in this study showed no differences in overall expression of NMDA receptors in multiple regions of the rodent cortex (Bird et al., 2015). Moreover, while PAE-dependent increases in the expression and sensitivity of GluN2B containing NMDARs in ventrolateral frontal cortex were observed, comparisons of sex and sex-by-alcohol interactions led to null results. These findings suggest perhaps other neurotransmitter systems or physiological characteristics such as brain vascularity, may underlie sex dependent changes in connectivity after moderate PAE.

The classification technique used in this study is sparsely utilized in animal neuroimaging studies. However, investigations utilizing machine learning to detect brain dysfunction in humans from fMRI FNC data are more established. For example, a study utilizing similar methods reported an accuracy measure of 84% in correctly detecting mTBI from a static FNC data set consisting of 48 patients and 48 healthy controls (Vergara et al., 2017). A follow up investigation that utilized dynamic FNC data found a 92% accuracy measure in detecting mTBI from one of several, yet unique, connectivity states (Vergara et al., 2018).

While the performance rate for detecting PAE is not as robust as those found in human studies of mTBI, it is important to consider a set of caveats. First, maternal blood alcohol levels during prenatal development reached a moderate 60.8 mg/dL (Davies et al., 2019). In rat studies of PAE, maternal alcohol serum levels can range from 30mg/dL (Cullen, Burne, Lavidis, & Moritz, 2014) in light exposure to 300 mg/dL (Mooney & Varlinskaya, 2011) in heavier exposure models. The rodent subjects from which FNC data were acquired were exposed to levels of prenatal alcohol on the low to moderate end of the range. Second, the alcohol-exposed offspring did not produce any detectable differences in brain volume nor blood perfusion in the frontal cortex when assessed in adulthood and compared to their respective control groups (Rodriguez, Davies, et al., 2016).

The results presented here, must also be considered within the context of a number of limitations. First, the FNC data utilized was of the static form which ignores temporal variations in connectivity across the scanning period. Examination of dynamic connectivity, which can account for these variations, may lead to disparate findings as evidenced in human-subjects research with dynamic FNC approaches showing better classification performance (Hutchison et al., 2013; Vergara et al., 2018). Future comparisons of dynamic and static FNC may reveal optimal approaches for the detection of PAE from FNC data. Furthermore, the neuroimaging data utilized to subsequently measure FNC was gathered from rodents under light isoflurane anesthesia. In our previous work, this approach was chosen to minimize the influence of motion during image acquisition. An alternative approach could employ the use of animal restraining devices to overcome anesthetic-related influences on brain function (King et al., 2005). In fact, studies conducted in rats and voles have revealed modest contributions of stress in normal and awake animals after an acclimation procedure (Liang, King, & Zhang, 2011, 2012; Reed, Pira, & Febo, 2013; Yee et al., 2016). However, changes in the sensitivity of stress-related circuitry including the hippocampus and the hypothalamic-pituitary-adrenal (HPA) axis following PAE are well documented (Hellemans, Verma, Yoon, Yu, & Weinberg, 2008; Lam, Raineki, Ellis, Yu, & Weinberg, 2018; Raineki, Ellis, & Weinberg, 2018), and carry the potential to introduce a different set of confounds in an awake scanning procedure. Next, animals in this investigation reached adulthood by the time image acquisition was conducted. Thus, additional research will need to examine machine learning detection in earlier postnatal developmental periods to enhance any potential translational utility of this approach. Our results are based off of a total sample size of 48, and a within-sex sample size of 24 (12 PAE; 12 SAC). Consequently, the procedures employed in this report stand to benefit from validation in additional contexts with increased samples to better leverage the utility of machine learning classifiers. Finally, the fMRI data were acquired in a 3.7T scanner. Future work with higher field strength scanners (e.g. 7T or 9T) may improve signal-to-noise ratio leading to increased classification performance.

## CONCLUSION

In summary, the application of support vector machine learning algorithms led to modest classification performance in discriminating alcohol-exposed from control animals utilizing FNC measures derived from the application of GICA to rodent resting state fMRI data. To our knowledge, this is the first study to apply machine learning classification methods to FNC data within the context of PAE. Future developments and refinements of the technique may lead to translational utility in human studies and lead to the development of novel and non-invasive ways of detecting FASD.

## ACRONYMS

BOLD: Blood Oxygen Level Dependent
FAS: Fetal Alcohol Syndrom
FASD: Fetal Alcohol Spectrum Disorder
fMRI: function Magnetic Resonance Imaging
FNC: Functional Network Connectivity
FWHM: Full Width Half Max
GICA: Group Independent Components Analysis
GIFT: Group ICA of fMRI Toolbox
HPA: Hypothalamic Pituitary Adrenal Axis
LOOCV: Leave One Out Cross Validation
MLP: Multilayer Perceptron
mTBI: mild Traumatic Brain Injury
NMDA: N-methyl-D-Aspartate
PAE: Prenatal Alcohol Exposure
QSVM: Quadratic SVM
RBF: Radial Basis Function
RSN: Resting State Network
SAC: Saccharin
SVM: Support Vector Machine

## Author Contribution Statement

C.R., V.V, V.C. and D.H. conceived the presented ideas. C.R., S.D., D.S., D.H., and V.C. contributed to the previously acquired data and preprocessing. C.R., V.V., performed additional preprocessing and data analyses. Manuscript writing efforts led by C.R. and V.V. with critical feedback from all authors.

